# Embryonic requirements for *Tcf12* in the development of the mouse coronal suture

**DOI:** 10.1101/2021.03.01.433456

**Authors:** Man-chun Ting, D’Juan T. Farmer, Camilla S. Teng, Jinzhi He, Yang Chai, J. Gage Crump, Robert E. Maxson

## Abstract

A major feature of Saethre-Chotzen syndrome is coronal craniosynostosis, the fusion of the frontal and parietal bones at the coronal suture. It is caused by heterozygous loss-of-function mutations in the basic HLH transcription factors *TWIST1* and *TCF12*. While compound heterozygous *Tcf12; Twist1* mice display severe coronal synostosis, the individual role of *Tcf12* has remained unexplored. Here we show that Tcf12 controls several key processes in calvarial development, including the rate of frontal and parietal bone growth, and the boundary between sutural and osteogenic cells. Genetic analysis supports an embryonic requirement for *Tcf12* in suture formation, as combined deletion of *Tcf12* in the embryonic neural crest and mesoderm, but not in the postnatal suture mesenchyme, disrupts the coronal suture. We also detect asymmetric distribution of Grem1 + mesenchymal cells on opposing sides of the wild-type frontal and parietal bones, which prefigures later bone overlap at the sutures. In *Tcf12* mutants, reduced asymmetry correlates with lack of bone overlap. Our results indicate a largely embryonic function of Tcf12 in controlling the rate and asymmetrical growth of calvarial bones and establishment of suture boundaries, which together ensure the proper formation of the overlapping coronal suture.

## Introduction

Craniosynostosis, the pathological fusion of the flat bones forming the top of the skull, occurs in approximately 1/2000 live births (Lajeunie et al., 1995). Fusion occurs at fibrous joints called sutures which allow the skull to flex during childbirth and mastication. Craniosynostosis occurs in both sporadic and syndromic forms, and often affects particular sutures (Wilkie et al., 2006; Johnson and Wilkie., 2011). The coronal suture forms between the frontal and parietal bones and is preferentially affected in Saethre-Chotzen syndrome. Heterozygous loss-of-function mutations in the basic helixloop-helix (bHLH) transcription factor genes *TWIST1* and *TCF12* account for most Saethre-Chotzen syndrome cases (Howard et al., 1997; El Ghouzzi et al., 1997; Sharma et al., 2013). Twist1 belongs to a subfamily of bHLH proteins whose members are expressed in specific tissues or cell types (Massari and Murre, 2000; Wang et al., 2015). Tcf12 is a member of the E protein subfamily of bHLH proteins, also including Tcf3 and Tcf4, whose members typically exhibit broad expression (Wang et al., 2015). In mice, heterozygous deletion of *Twist1* phenocopies the selective coronal suture loss seen in human Saethre-Chotzen patients (El Ghouzzi et al., 1997). Whereas heterozygosity of *Tcf12* alone does not result in suture defects in mice, it does enhance coronal synostosis in *Twist1+/−* mice (Sharma et al.,2013). In zebrafish, loss of *tcf12* or *twist1b* alone does not cause craniosynostosis, yet combined homozygous loss of *twist1b* and *tcf12* results in coronal-specific synostosis (Teng et al., 2018), despite the zebrafish coronal suture likely being non-homologous with the murine coronal suture (Teng et al., 2019). However, the individual role of Tcf12 in skull development has remained unexplored in model organisms, leaving it unclear whether Twist1 and Tcf12 control similar or distinct processes during suture development.

In mice and zebrafish, combinatorial loss of *Twist1* and *Tcf12* alleles influences the dynamics of osteogenic precursors during embryonic calvarial bone development (Teng et al., 2018). In compound mutants, frontal and parietal bone growth was accelerated, with the degree of acceleration in zebrafish predictive of later coronal suture loss. In addition to effects on osteogenic precursor dynamics in compound *Twist1; Tcf12* heterozygous mice, heterozygous loss of *Twist1* also results in inappropriate cell mixing at the boundary between the neural crest-derived frontal and mesoderm-derived parietal bone (Merrill et al., 2006; Ting et al., 2009). *Twist1* function is required in both neural crest-derived and mesoderm-derived cells for suture formation, with only heterozygous loss of *Twist1* in both tissues resulting in synostosis (Teng et al., 2018). However, as these studies used non-conditional Cre drivers, distinct requirements in embryonic versus postnatal sutures could not be distinguished. This is a significant caveat as recent work has demonstrated that the postnatal suture is a continuous source of osteogenic cells for the growth and repair of the calvarial bones. For example, *Gli1:CreER* can be used to mark long-term stem cells in the postnatal mouse suture, with genetic ablation of these labeled cells resulting in loss of all sutures (Zhao et al., 2015). Whereas *Twist1* and *Tcf12* are expressed in postnatal suture mesenchyme, we show that conditional deletion of *Tcf12* in the postnatal *Gli1+* sutural stem cell domain does not affect maintenance of sutures, pointing to largely embryonic roles of Twist1 and Tcf12 in suture regulation.

Twist1 can form homodimers as well as heterodimers wirh a variety of bHLH factors, including Tcf12 and Tcf3 (Fan et al., 2020). Forced dimer approaches in ES cells (Fan et al., 2020) and in mouse sutures (Connerney et al., 2006) provided evidence that Twist1 homodimers and Twist1-E heterodimers drive different cellular processes. In differentiating ES cells, heterodimers promote differentiation of ectodermal and mesodermal markers and homodomers maintain cells in a pluripotent state. In developing mouse sutures, a higher Twist1/Twist1 to Twist1/E ratio was associated with an expansion of the osteogenic fronts and suture closure. It thus seems clear that a change in the ratio of Twist1 homodimers versus Twist1-E heterodimers can produce different biological outcomes, although the exact role of Twist1 homodimers and heterodimers in suture development and craniosynostosis remains unclear.

Here we investigate the extent to which Tcf12, a binding partner for Twist1 (Fan et al., 2020), participates in similar developmental processes of suture formation as described for Twist1. In contrast to the expectation from forced dimer approaches, we show that homozygous loss of *Tcf12* recapitulates many of the calvarial phenotypes of heterozygous *Twist1* and compound heterozygous *Twist1; Tcf12* mice, including accelerated frontal and parietal bone growth, and disruption of the boundary between sutural and osteogenic cells. We also find that Tcf12 function is required in both neural crest-derived and mesoderm-derived cells for coronal suture formation. Interestingly, we find that Tcf12 also controls the asymmetry of Grem1+ mesenchyme cells at the parietal and frontal bone fronts, which may help explain the predictable overlapping nature of these bones at the normal coronal suture.

## Materials and Methods

### Mouse mutants and genotyping

The *Tcf12* (Wojciechowski et al.,2007), *Twist1* (Chen and Behringer, 1995), *R26R* (Soriano., 1999), *Wnt1-cre* (Daniellian et al.,1998) and *Mesp1-cre* (Saga et al., 1999) alleles have been described. We genotyped *Tcf12, Twist1, R26R, Wnt1-cre* and *Mesp1-cre* alleles by PCR as described (Wojciechowski et al.,2007; Chen and Behringer, 1995; Saga et al., 1999; Jiang et al., 2002; Teng et al., 2018)

### Whole mount skull alizarin red S staining

Skulls of late-stage embryos, new born (P0), and post-natal mice were stained for bone with 2% Alizarin Red S in 1% KOH for 3 to 5 days. The specimens were then cleared and stored in 100% glycerol.

### Whole mount alkaline phosphatase (ALP) and LacZ staining

E13.5 and E14.5 heads were fixed in 4% paraformaldehyde (PFA) in PBS, and were bisected midsagitally after fixation. Presumptive calvarial bones were stained with NBT and BCIP (Roche). To detect LacZ, skulls of late-stage embryos were fixed in cold 4% PFA for 30 minutes followed by PBS washes. Calvarial bones were then stained with b-galactosidase staining solution overnight at 37°C.

### Histology and immunohistochemistry

To evaluate cellular proliferation at E14.5, BrdU (Sigma) was injected into the pregnant female (200 μg/g body weight) 2 hour prior to dissection. Heads of embryos were embedded in OCT medium (Histoprep, Fisher Scientific) before sectioning. Frozen sections were cut at 10 μm. Immunohistochemistry was performed using rat anti-BrdU (MCA2060 GA, Bio-Rad), rabbit anti-Sp7/Osx (sc-22536-r, Santa Cruz), rabbit anti-ephrin-A2 (sc-912, Santa Cruz), and goat anti-Grem1 (PA5-47973, ThermoFisher) diluted in 1% BSA/PBS and incubated overnight at 4°C. Detection of primary antibody of anti-Brdu, anti-Osx, anti-ephrinA2, and anti-Grem1 was performed by incubating goat anti-rat-FITC (sc-2011, Santa Cruz), goat anti-rabbit-Alexa Fluor 568 (ab1175471, abcam), donkey anti-rabbit-Alexa Fluor 568 (ab1175470, abcam), and donkey anti-goat IgG-Alexa Fluor 488 (ab150129, abcam) respectively for one hour at room temperature followed by DAPI counterstaining and examination by epifluorescence microscopy. Analysis of β-galactosidase activity of *Wnt1-Cre/R26R* reporter gene expression was carried out as described (Merrill et al., 2006). Postnatal calvaria were dissected and fixed in 10% neutralized buffered formalin (NBF, Sigma) overnight at room temperature, then decalcified with 10% EDTA in 1xPBS for 1-14 days based on mouse age. Decalcified calvaria were dehydrated and embedded in paraffin. Tissue blocks were sectioned at 5μm using a microtome (Leica) and mounted on SuperFrost Plus slides (Fisher). Hematoxylin and Eosin (H&E) staining was completed following standard protocols.

### RNAscope

RNAscope in situ hybridization for embryonic timepoints was performed with the RNAscope Multiplex Fluorescent Kit v2 (Advanced Cell Diagnostics, Newark, CA) according to the manufacturer’s protocol for the fixed-frozen sections. Heat antigen retrieval was omitted to maximize nuclei quality, and TSA^®^ Plus (Fluorescein, Cy3 and Cy5) reagents were used at 1:1000 dilution. For postnatal calvaria, tissues were fixed in 10% NBF overnight at room temperature, decalcified with DEPC-treated 10% EDTA in 1×PBS at 4°C, dehydrated sequentially with 15% sucrose in 1xPBS (4°C, overnight) and 30% sucrose in 1xPBS/OCT (4°C, overnight), and embedded in OCT (Sakura, Tissue-Tek, 4583) on liquid nitrogen immediately. Frozen tissue blocks were sectioned at 8μm on a cryostat (Leica CM3050S) and mounted on SuperFrost Plus slides. Probes for *Tcf12* and *Twist1* were purchased from Advanced Cell Diagnostics (Cat #. 504861). In situ RNA detection was performed using RNAscope^®^ 2.5 HD Detection Reagent (Advanced Cell Diagnostics, 322360) according to instructions provided by the manufacturer.

### Quantitation and statistical analyses

The severity of coronal synostosis was quantified using a scoring method adapted from Oram and Gridley’s craniosynostosis index (Oram and Gridley, 2005). For assessment of BrdU+, Sp7+ and Grem1+ cells in mouse coronal suture area, stained-positive cells in a defined area across the bone fronts and suture were manually counted. Five sections per animal were quantified and averaged. Pairwise comparisons were analyzed by two-tailed Student’s t-test, and the p-values were corrected for multiple testing with Bonferroni correction.

## Results

### Homozygous loss of *Tcf12* disrupts coronal suture formation

To investigate the role of the murine *Tcf12* gene in the development of craniosynostosis, we generated a *Tcf12* null allele by crossing *Tcf12^flox/flox^* mice with EIIa-Cre, a general deleter active in the germline (Lakso et al., 1996). From a *Tcf12^+/−^* incross, homozygous *Tcf12^−/−^* mice were recovered at low frequency, with the majority of surviving pups dying within the first 2 weeks, consistent with previous findings (Zhuang et al., 1996). In order to examine the morphology of the coronal suture, we carried out Alizarin Red staining on the skulls from homozygous *Tcf12* mutants surviving into postnatal stages (P0-P21). We found partial fusion of the coronal suture in 8 of 22 *Tcf12^−/−^* mice (36%), and no coronal suture defects in 17 heterozygous *Tcf12^+/−^* and 15 wild-type sibling mice (Fig 1 and Table 1). We also observed developmental delay and rare exencephaly in *Tcf12^−/−^* mice, as reported by (Zhuang, et al. 1996), with the frequency of exencephaly increasing from 3% to 33% when one allele of *Twist1* was also deleted (Table 1). *Tcf12^−/−^* mice also displayed kinks and curls of the tail, a phenotype known to be caused by a moderate delay in posterior neuropore closure (van Straaten and Copp, 2001). *Tcf12^−/−^*; *Twist1^+/−^* mice also infrequently displayed an open ventral body wall not observed in *Tcf12^−/−^* mice (Table 1). The suture phenotypes and low penetrance exencephaly and curled tail defects suggest *Tcf12* functions in both calvarial suture development and neurulation.

**Figure 1.**
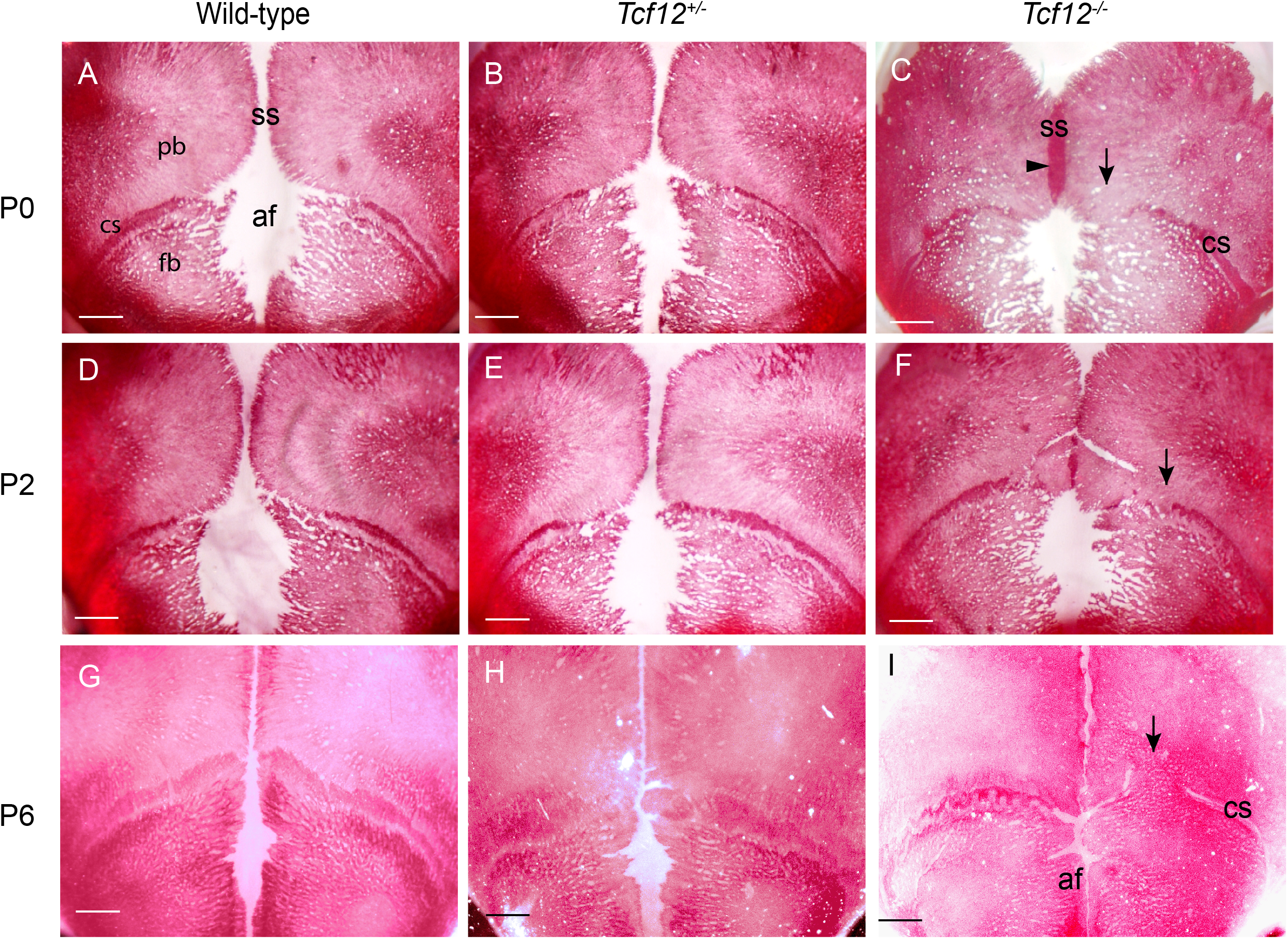
Fusion of coronal sutures in *Tcf12* null mice. Skulls of mice were stained with Alizarine Red S. Note open coronal sutures (cs) in wild-type (A, D, G) and *Tcf12^+/−^* mice (B, E, H) versus unilateral partially fused coronal sutures (C, F, I, arrows) and narrowed sagittal suture (C, ss, arrowhead) in *Tcf12^−/−^* null mice. P0, P2, P6: postnatal days 0, 2, and 6. Scale bars = 1 mm.

**Table 1.** Phenotypes of *Tcf12* mutants

| Mouse Phenotypes<br>(n) | *Tcf12*<sup>-/-</sup><br>(32) | *Twist*<sup>+/-</sup>;*Tcf12*<sup>+/-</sup><br>(46) | *Twist*<sup>+/-</sup>;*Tcf12*<sup>-/-</sup><br>(21) |
| --- | --- | --- | --- |
| Coronal Craniosynostosis | 58% | 100% | 100% |
| Curly Tail | 12.5% | 0% | 19% |
| Malocclusion | No | Yes | N/A |
| Exencephaly | 3% | 0% | 33% |
| Developmental delay | Yes | Yes | Yes |
| Open Ventral Body Wall | 0% | 0% | 14% |

Mice of the indicated genotypes were examined within the first postnatal week. To assess coronal craniosynostosis, skulls were stained with Alizarin Red S and scored according to the method of Oram and Gridley (2005).

**Table 2.** Severity of craniosynostosis in different genetic combinations

| Genotype | n | Coronal<br>Craniosynostosis index<br>(C.I.) ± SEM | Penetrance<br>% |
| --- | --- | --- | --- |
| *Wildtype* (P21) | 15 | 0.0 ± 0.0 | 0 |
| *Tcf12*<sup>+/-</sup> (P21) | 17 | 0.0 ± 0.0 | 0 |
| *Tcf12*<sup>-/-</sup> (P0-P21) | 22 | 0.5 ± 0.2 | 36 |
| *Tcf12*<sup>flox/flox</sup>; *Wnt1-Cre*; *Mesp1-Cre*<br>(P0-P21) | 6 | 1.7 ± 1.8 | 66.7 |
| *Twist1*<sup>+/-</sup> | 9 | 2.2 ± 0.5 | 100 |
| *Tcf12*<sup>+/-</sup>; *Twist1*<sup>+/-</sup> | 6 | 4.8 ± 0.4 | 100 |

Calvaria were dissected at the indicated developmental stages and stained with Alizarin Red S. The degree of fusion of the coronal suture was assessed as described by Oram and Gridley (2005).

### Tissue-specific requirements of *Tcf12* for suture patency and calvarial bone growth

To investigate where *Tcf12* is required for suture development and skull bone growth, we deleted both copies of *Tcf12* in neural crest (*Tcf12^flox/flox^*; *Wnt1-Cre*), mesoderm (*Tcf12^flox/flox^*; *Mesp1-Cre*), or both (*Tcf12^flox/flox^*; *Wnt1-Cre; Mesp1-Cre*). Whereas homozygous loss of *Tcf12* in either neural crest or mesoderm alone did not affect coronal suture patency, loss in both tissues prevented coronal suture formation (Fig 2A-H). The severity of coronal suture defects was higher in combined conditional mutants (coronal synostosis index 1.7 +/− 1.8) than conventional homozygous *Tcf12* mice (coronal synostosis index 0.5 +/− 0.2), perhaps reflecting the prenatal lethality of more severely impacted conventional null mice. Given a similar requirement for *Twist1* in both mesoderm- and neural crest-derived tissues (Teng et al., 2018), these results support *Tcf12* and *Twist1* acting together in the same tissues for coronal suture formation.

**Figure 2.**
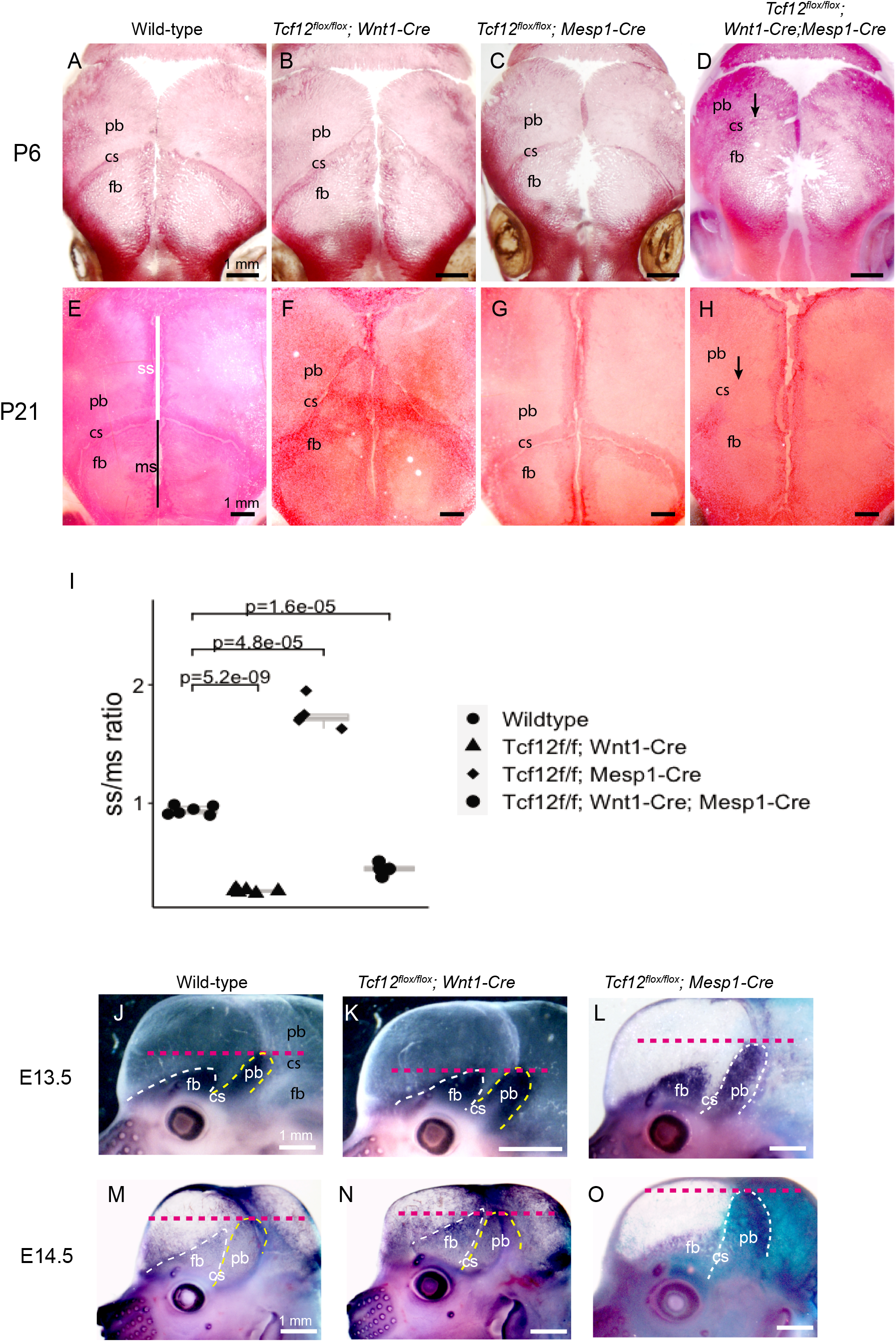
Tissue-specific *Tcf12* requirements in calvarial bone growth and suture patency. (A-H) Skulls of the indicated genotypes were stained with Alizarin Red S at postnatal day 6 (P6) and 21 (P21). Note the change in the relative size of the frontal and parietal bones in *Tcf12^flox/flox^, Wnt1-Cre and Tcf12^flox/flox^, Mesp1-Cre* mutants. Loss of the coronal suture (arrows, often bilateral) was only observed upon deletion of *Tcf12* from both the neural crest and mesoderm *(Tcf12^flox/flox^, Wnt1-Cre; Mesp1-Cre)*. (I) Quantification of the ratio of the sagittal suture (ss) to the metopic suture (ms) illustrates relative expansion of the frontal bone (low ss/ms ratio) and parietal bone (high ss/ms ratio) upon deletion of *Tcf12* from the neural crest or mesoderm lineages, respectively. P values of pairwise comparisons were obtained from a twotailed Student’s t-test. Error bars represent standard error of the mean. (J-O) Lateral views of Alkaline Phosphatase (ALP) staining of frontal and parietal bone rudiments (white dashed lines). Note apical expansion of the frontal bone rudiment in *Tcf12^flox/flox^;Wnt1-Cre* embryos, and apical expansion of the parietal bone rudiment *in Tcf12^flox/flox^; Mesp1-Cre* embryos. The red dotted lines indicate the extent of the expansion of the frontal or parietal bone primordia. The embryo shown in O carries the R26R allele and is stained for lacZ. fb: frontal bone; pb: parietal bone; cs: coronal suture; ss: sagittal suture; ms: metopic suture. Scale bars = 1 mm.

In both *Tcf12* null mice and those with conditional homozygous *Tcf12* deletion in the mesoderm, we observed a narrowing of the sagittal suture at P6, consistent with accelerated growth of the mesoderm-derived parietal bones (Fig 1, Fig 2C). Upon mesodermal deletion, we also observed an expansion of the size of the parietal bone relative to the neural crest-derived frontal bone at P21, effectively moving the coronal suture anteriorly (Fig 2G, I). Reciprocally, the size of the frontal bone expanded relative to the parietal bone upon neural crest deletion of *Tcf12*, shifting the coronal suture posteriorly (Fig 2F, I). This relative size change was associated with increased apical growth of the frontal or parietal bone rudiments at E13.5 and E14.5 in *Tcf12^flox/flox^; Wnt1-Cre* or *Tcf12^flox/flox^; Mesp1-Cre* mutants, respectively, as revealed by alkaline phosphatase (ALP) staining (Fig 2J-O).

At E18.5 and P1, the apical expansion of the frontal bones in *Tcf12^flox/flox^; Wnt1-Cre* mice was associated with the appearance of Alizarin-Red positive heterotopic bones in the normally unossified area when compared with controls (Fig 3A-F). Sectioning and whole-mount analysis of neural crest-derived mesenchyme (*Tcf12^flox/flox^; Wnt1-Cre; R26RLacZ*) at E15.5 and P1 showed that the ectopic bone is entirely of neural crest origin (Fig 3F-N). Thus, the expansion of the frontal relative to the parietal bone upon neural crest-specific *Tcf12* loss is due, at least in part, to accelerated osteoblast differentiation within neural crest-derived cells.

**Figure 3.**
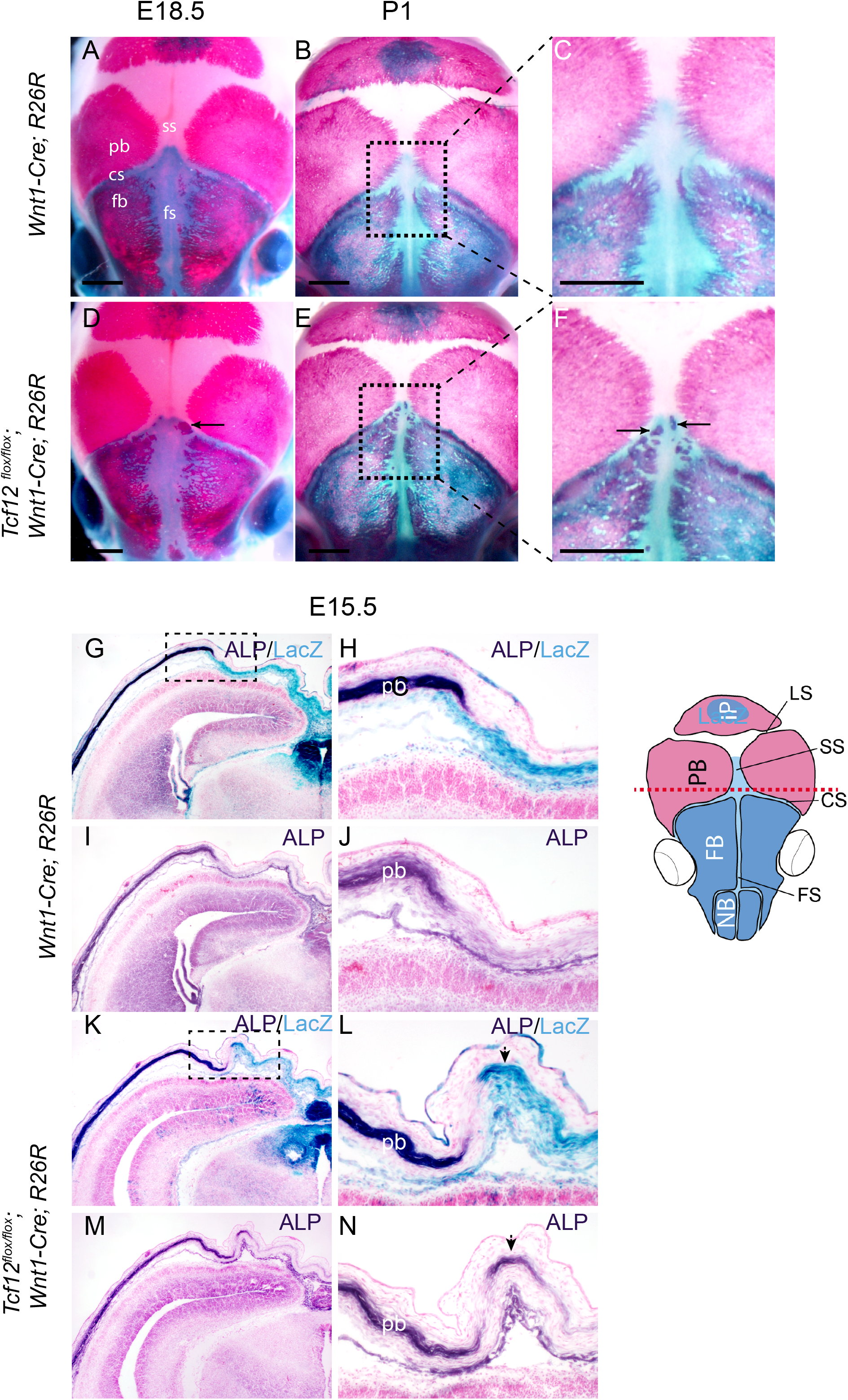
Accelerated frontal bone formation upon neural crest deletion of *Tcf12*. (A-F) Skulls of the indicated genotypes were stained with Alizarin Red S (bone) and for lacZ (blue, neural crest-derived cells) at E18.5 (A, D) or P1 (B, C, E, F). Note bony islands (arrows) in lacZ-positive neural crest-derived cells situated in the anterior sagittal suture of *Tcf12^flox/flox^; Wnt1-Cre; R26R* embryos (D, magnified region in F). (G-N) Adjacent cross sections of embryos of the indicated genotypes were stained for alkaline phosphatase (purple, osteoblasts) or lacZ (blue, neural crest-derived cells) at E15.5. Boxed areas correspond to enlarged regions on the right. Note the bony islands in adjacent sections shown in L and N are lacZ-positive. Diagram of the E15.5 mouse skull on the right shows approximate location of sections. Red is mesoderm-derived and blue is neural crest-derived tissue. cs, coronal suture; fb, frontal bone; fs, frontal (metopic) suture; pb, parietal bone; ss, sagittal suture; ip, interparietal bone; ls, lambdoid suture, nb, nasal bone. Scale bars = 1 mm.

### Reduction of *Tcf12* function affects osteoblast dynamics and asymmetric distribution of Grem1+ mesenchyme

We showed previously that closer apposition of frontal and parietal bones in *Twist1^+/−^: Tcf12^+/−^* mutants was associated with increased proliferation yet reduced number of putative progenitors at the growing bone fronts and the developing coronal suture (Teng et. al., 2018). Given the accelerated frontal and parietal bone growth we observed upon *Tcf12* deletion, we assessed whether osteoblast and progenitor dynamics were also altered in *Tcf12* null mice. At E14.5, we observed an approximately 22% increase in the number of Sp7+ osteoblasts along the frontal and parietal bone fronts of *Tcf12* mutants, yet the percentage of Sp7+ osteoblasts incorporating BrdU (a marker of cell proliferation) and the total number of proliferative Sp7-cells were unchanged (Figure S1). These findings indicate that both *Tcf12* and *Twist1* function to inhibit premature osteoblast differentiation, though only Twist1 appears to restrict proliferation at the bone fronts.

We next examined the number of putative osteoprogenitors at the developing coronal suture in *Tcf12* mutants. Grem1 marks mesenchymal progenitors responsible for bone formation in the appendicular skeleton (Worthley et al., 2015), and we had previously shown that the number of Grem1-positive cells around the frontal and parietal bones during coronal suture formation was reduced in *Tcf12^+/−^; Twist1^+/−^* mice (Teng et. al., 2018). Interestingly, we observed that Grem1 + cells were asymmetrically distributed above (lateral to) the frontal bone and below (medial to) the parietal bone in wild-type E14.5 controls (Fig 4D, F, H and I). This asymmetric distribution of Grem1+ cells correlates with the parietal bone reproducibly overlapping above the frontal bone at the mature coronal suture. In *Tcf12* null mice, Grem1+ cells above the frontal bone were selectively lost (Fig 4E, G and H), resulting in a more symmetric arrangement of Grem1+ cells. Grem1+ cells both above and below the parietal bone were also reduced in mutants. Consistently, we observed that *Tcf12* and *Twist1* expression was asymmetrically enriched in mesenchyme above the frontal bone at E15.5 (Fig 4A-C, arrows). These results indicate a requirement of *Tcf12* for the asymmetric enrichment of Grem1+ mesenchymal cells above the frontal bone, with loss of asymmetry in mutants correlating with the frontal and parietal bones meeting end-on-end and often fusing.

**Figure 4.**
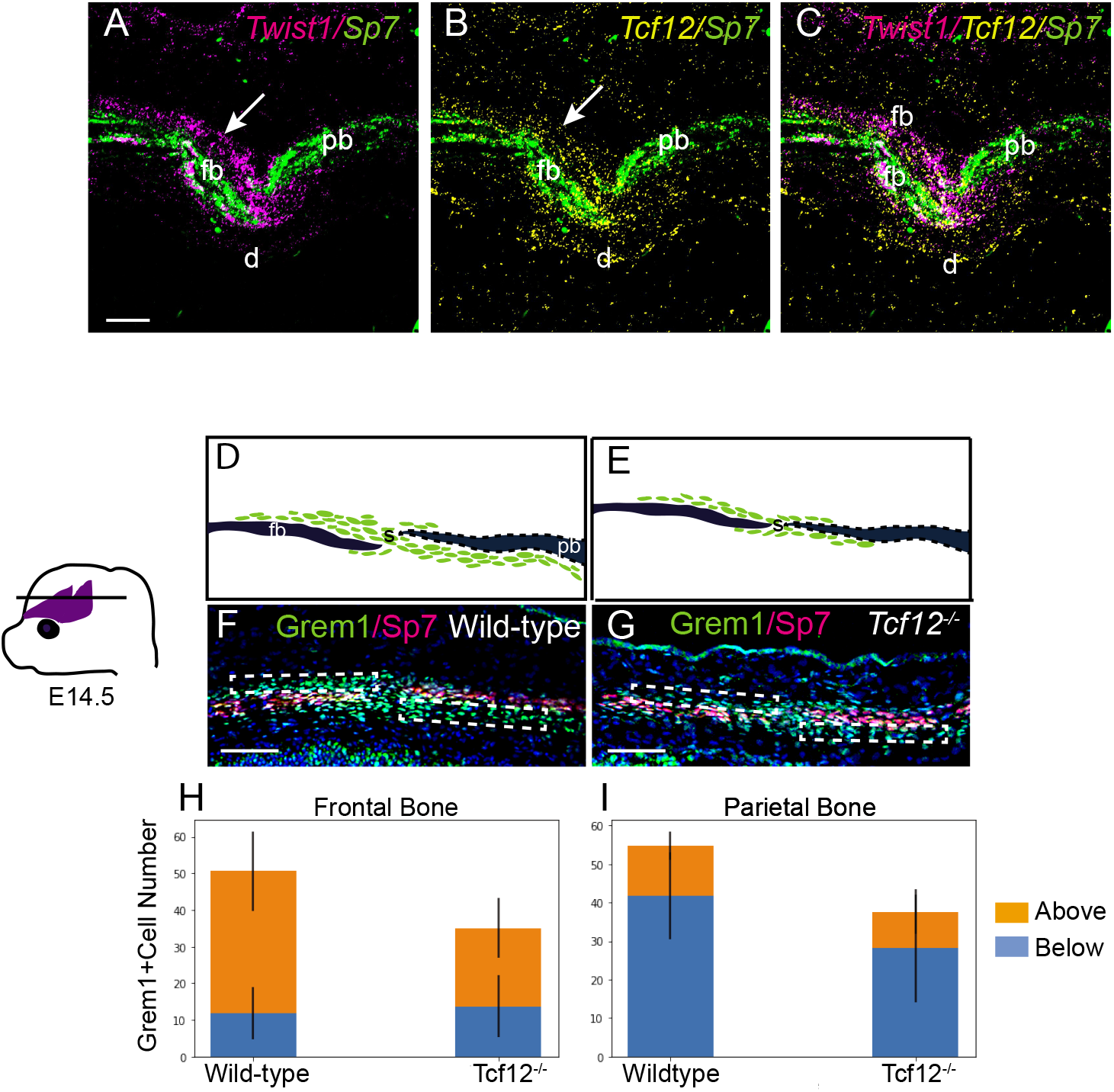
Asymmetric distribution of Grem1+ mesenchyme around the frontal and parietal bones at the coronal suture. (A-C) RNAscope in situ hybridization of the coronal suture region at E14.5, with arrows showing asymmetric distribution of mesenchyme expressing *Twist1* and *Tcf12* above the frontal bone (labeled by *Sp7* expression in green). (D-G) Immunostaining of sections through the coronal sutures of E14.5 embryos with antibodies against Grem1, a putative marker of osteoprogenitor cells, and Sp7, a marker of osteoblasts. Diagrams of the bones (black), and suture and bone front mesencnyme (green), are shown above. (H, I) Quantification of the Grem1+ cells in the boxed regions in F and G. Error bars represent the standard error of the mean. P values: above vs below the wild-type frontal, p = 1.3×10^−12^; above vs below the wild-type parietal, p = 9.5×10^−15^; above the wild-type frontal vs above the mutant frontal, p = 2.3×10^−7^; below the wild type frontal vs below the mutant frontal, p = 0.42; above the wild-type parietal vs above the mutant parietal, p = 0.02; below the wild-type parietal vs below the mutant parietal, p = 8.0×10^−4^. Scale bars = 50 μm.

### Neural crest-mesoderm boundary defects in *Tcf12* null mice

Disruption of the boundary between the osteogenic neural crest cells of the frontal bone and the mesoderm-derived coronal suture mesenchyme was observed in *Twist1* heterozygous mice and was accompanied by a reduction in Eph-ephrin signaling in the ectocranial layer above the coronal suture (Merrill et al., 2006). We used *Wnt1-Cre; R26R* mice to trace the fate of neural crest-derived cells at the coronal sutures of E13.5 embryos. In wild types, LacZ-positive neural crest-derived cells contributed to the frontal bone and were excluded from the mesoderm-derived coronal suture mesenchyme and parietal bone. In contrast, neural crest-derived cells invaded the coronal suture mesenchyme in *Tcf12^−/−^* mutants, consistent with a defect in the neural crest-mesoderm boundary (Fig 5A-F). We next examined ephrinA2-EphA signaling in *Tcf12* null mice, as we had previously shown that this pathway is required in the ectocranial layer above the forming suture to regulate neural crest-derived cell migration and boundary formation (Merrill et al., 2006; Ting et al., 2009; Yen et al., 2010). In *Tcf12^−/−^* mutants, ectocranial expression of *Efna2* was completely abolished (Fig 5G-J). We conclude that Tcf12 and Twist1 act similarly to regulate *Efna2* ectocranial expression and the boundary between osteogenic and non-osteogenic cells at the nascent coronal suture.

**Figure 5.**
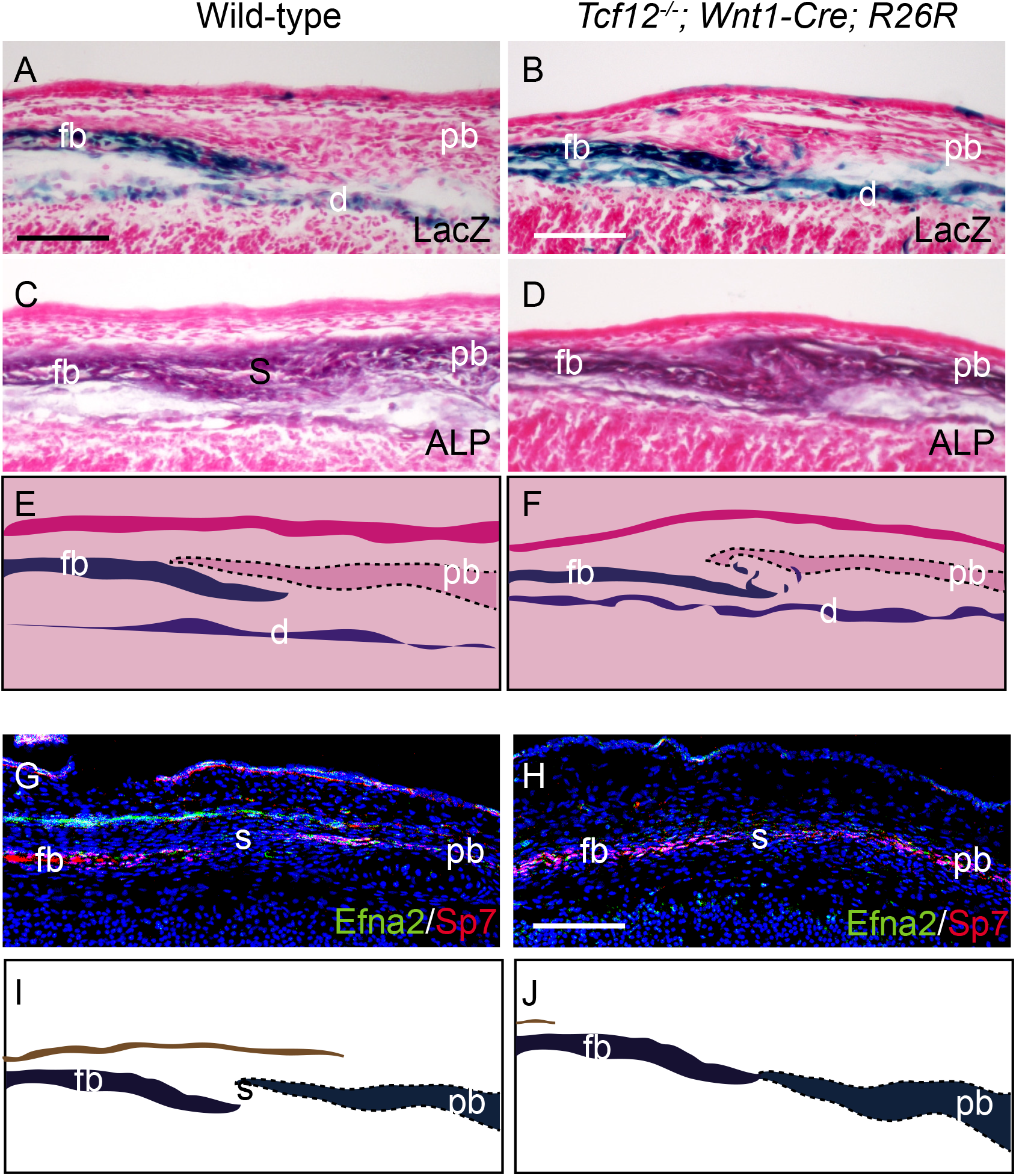
Disruption of the neural crest-mesoderm boundary at the *Tcf12* mutant coronal suture. Sections of wild-type *Wnt1-Cre; R26R* and *Tcf12^−/−^; Wnt1-Cre; R26R* embryos at the level of the coronal suture were stained for lacZ (A, B), alkaline phosphatase (ALP) (C, D), and EfnA2 and Sp7 (G, H) at E13.5. In *Tcf12^−/−^* mutants, *lacZ*-positive neural crest cells are located ectopically in the area of the prospective coronal suture (B, diagrammed in F), and Efna2 is lost in the ectocranial layer (H, diagrammed in J). fb: frontal bone; pb: parietal bone; s: coronal suture. Scale bars = 50 μm

### *Tcf12* is dispensable in postnatal suture mesenchyme for suture patency

We next investigated potential roles for Tcf12 in regulating suture patency at postnatal stages. Using RNAscope in situ hybridization of 14-week-old mouse skulls, we observed postnatal expression of *Tcf12* throughout the mesenchyme of the coronal suture, as well as in periosteum above and dura mesenchyme below the suture (Fig 6A). As the suture mesenchyme expression of *Tcf12* overlaps with labeling by *Gli1-CreERT2* at 4 weeks after birth, we treated *Gli1-CreERT2; Tcf12^flox/flox^* mice at this stage with tamoxifen to delete *Tcf12* in the postnatal suture mesenchyme. *Gli1-CreERT2* has been used to inactivate *Twist1* in adult mice (Yu et al., 2021). Fourteen weeks after induction, histology revealed no defects in the coronal suture when compared with untreated wild-type controls (Fig 6B,C). This finding argues against a critical postnatal role of *Tcf12* in the suture mesenchyme for maintaining suture patency, although we cannot rule out postnatal roles for *Tcf12* in the periosteum and dura.

**Figure 6.**
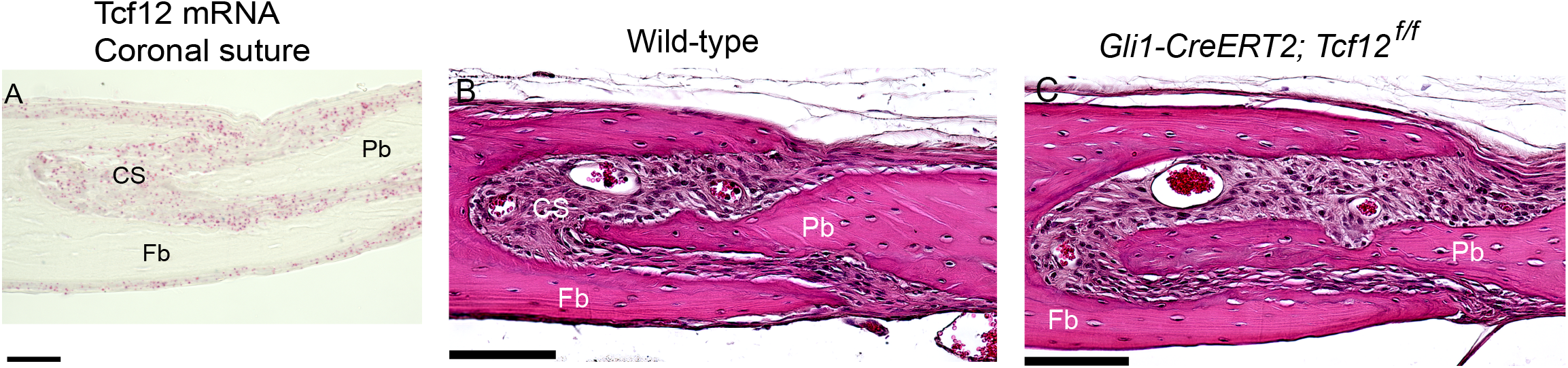
Postnatal deletion of *Tcf12* in suture mesenchyme does not affect coronal suture maintenance. (A) RNAscope in situ hybridization on sections of 14-week postnatal mice shows expression of *Tcf12* (red dots) in the coronal suture mesenchyme, as well as periosteum (above bones) and dura (below bones). (B, C) Sections through the coronal suture of a wild type and a *Tcf12^flox/flox^; Gli1-CreERT2* mouse stained with Aliazarin Red and hematoxylin/eosin. Tamoxifen was administered at 4 weeks after birth and skulls were analyzed at 14 weeks. No defects in coronal suture maintenance were observed. CS, Coronal Suture; Fb, Frontal bone; Pb, Parietal bone. Scale bars = 50 μm.

## Discussion

The ability of heterozygous mutations in *TWIST1* or *TCF12* to cause coronal synostosis in Saethre-Chotzen syndrome, as well as genetic interactions of *Twist1* and *Tcf12* homologs in zebrafish and mice, had suggested that *Twist1* and *Tcf12* may similarly control several key processes in calvarial development. However, *Twist1* can also form homodimers, as well as heterodimers with other partners besides *Tcf12*, suggesting that there could be differences between Twist1 and Tcf12 roles during coronal suture development. Here we show that *Twist1^+/−^* and *Tcf12^−/−^* mutants share many but not all defects related to coronal suture formation, suggesting a primary role for Twist1-Tcf12 heterodimers in this process.

We find that *Tcf12^−/−^* mutants display the same partially penetrant coronal synostosis as *Twist1^+/−^* mutants, which is similarly preceded by accelerated frontal and parietal bone growth and disruption of the boundary between sutural and osteogenic cells (Merrill et al., 2006; Yen et al., 2010; Ting et al., 2009; Teng et al., 2018). At a molecular level, this is reflected by an increase in Sp7+ osteoblasts and a concomitant decrease in Grem1+ mesenchyme cells at bone fronts, as well as a loss of ectocranial *Efna2* expression. As with *Twist1, Tcf12* must be mutated in both the neural crest and mesoderm lineages to disrupt the coronal suture. These similarities suggest that Tcf12 and Twist1 may act together to control common gene targets and cellular processes important for proper coronal suture development. We did not, however, observe the same increases in osteoblast and/or progenitor proliferation in *Tcf12* mutants as reported for *Twist1^+/−^* mutants (Teng et al., 2018). This difference could be explained by the milder suture fusion seen in *Tcf12^−/−^* mutants, and/or a selective requirement of Twist1 homodimers for restricting the proliferative expansion of bone-forming cells. Rather than controlling proliferation, our lineage analysis of neural crest-derived cells in *Tcf12* mutants suggests that Tcf12 functions primarily to restrict osteoblast differentiation at the growing bone fronts.

Whereas heterozygous loss-of-function mutations of *TCF12* in humans cause coronal synostosis, *Tcf12* heterozygosity does not in mice. Moreover, even *Tcf12* null mice displayed a weaker synostosis phenotype than *Twist1* heterozygous mice. One explanation is that the perinatal lethality of many *Tcf12* null mice selects against more severely affected mutants. Supporting this claim, conditionally deleting *Tcf12* in neural crest and mesoderm increases viability and the severity of coronal synostosis. Another possibility is that there is compensation by related E-box bHLH factors.

Our results suggest several potential mechanisms that may lead to synostosis upon loss of *Tcf12* and/or *Twist1*. Mutation of either gene results in excess osteoblast production and acceleration of frontal and parietal bone growth. This is clearly seen upon inactivation of *Tcf12* in the neural crest. In these animals, small disorganized islands of bone form in the frontal fontanelle, the space where the posterior portion of the parietal bones and anterior portion of the frontal bones come together. These bony islands arise from neural crest-derived cells of the frontal foramen, consistent with inappropriate conversion of non-osteogenic neural crest-derived cells to osteoblasts. Why only inactivation of *Tcf12* or *Twist1* in both the neural crest and mesoderm causes synostosis is unclear. In one model, stem cell depletion in one lineage can be compensated by stem cells from the other lineage. Hence, depletion in both lineages would be required to bring stem cell numbers below the threshold necessary to maintain separate bones. Consistent with this model, we observe depletion of Grem1+ mesenchyme cells associated with the bone fronts in both *Twist1^+/−^* and *Tcf12^−/−^* mutants, though future lineage tracing will be required to determine whether Grem1+ cells act as osteogenic stem cells as described in the limbs (Worthley et al., 2015). In addition to acceleration of bone growth, disruption of either *Tcf12* or *Twist1* also results in a disruption of the boundary between the frontal bone and the suture mesenchyme that is associated with loss of overlying *Efna2+* ectocranial expression. Whether this boundary defect is a secondary consequence of the accelerated bone growth, or is an independent phenotype that further contributes to synostsosis, requires further investigation.

Intriguingly, our results provide an explanation for the overlap of the frontal and parietal bones at the coronal suture. Unlike the medially placed interfrontal and sagittal sutures, which are formed by the direct apposition of osteogenic fronts on a level plane, the transversely situated coronal suture is reproducibly formed by the parietal bone overlapping above the frontal bone. In wild types, we find a preferential enrichment of Grem1+ mesenchyme cells above the frontal bone and below the parietal bone, with *Tcf12* and *Twist1* showing similar asymmetric expression, in particular above the frontal bone rudiment. One possibility is that these asymmetries skew the growth of the two bones off their central axes such that that the parietal reproducibly overlaps above the frontal bone. In mice deficient for *Tcf12* or *Twist1*, the osteogenic fronts of the frontal and parietal bones fail to overlap, and this correlates with a more symmetrical distribution of Grem1+ cells in relation to the growing frontal bone. In contrast, the asymmetric distribution around the parietal bone was less affected. One explanation for the reduction in frontal but not parietal bone mesenchymal asymmetry is loss of the *Efna2+* ectocranial layer in mutants. The ectocranial *Efna2+* layer overlays the enriched Grem1 + cells of the frontal bone but does not significantly extend into the parietal side of the coronal suture. It is therefore possible that the ectocranial layer serves to enrich Grem1+ cells above the frontal bone in normal animals, with loss of this ectocranial layer in mutants disrupting asymmetry and leading to increased apposition of bone fronts that could further contribute to synostosis.

---

**Figure S1. Osteogenic cell dynamics in the *Tc12^−/−^* coronal suture.** (A, B) Sections of E14.5 embryonic heads stained with antibodies against Sp7 and BrdU, after injecting BrdU into pregnant females two hours prior to harvesting embryos. Counts were performed in the boxed areas, which include the osteogenic fronts of the frontal and parietal bones and the interposed mesenchyme. (C-E) Quantification of Sp7+ osteoblasts, proliferative Sp7−/BrdU+ cells, and proliferative Sp7+/BrdU+ in the boxed regions in A and B. Error bars represent the standard error of the mean. P values: Sp7+ osteoblasts, mutant vs wild-type, p = 0.016, proliferative Sp7−/BrdU+ cells, mutant vs wild-type, p = 0.75; proliferative Sp7+/BrdU+ cells, p = 0.31. Scale bars = 50 μm.

## Acknowledgements

This work was supported by NIH grant R01DE026339 to REM, JGC, and YC, and NIH grant R35DE027550 to JGC. CST was supported by NIH training fellowships F31DE024031 and T90DE021982, and DTF was supported by the HHMI Hanna H. Gray Fellows Program. We thank members of the Maxson, Crump and Chai labs for helpful discussions.

